# Decreased KCC2 expression in the human spinal dorsal horn associated with chronic pain and long-term opioid use

**DOI:** 10.1101/2025.01.19.633733

**Authors:** Olivia C Davis, Samuel Ferland, Louis-Etienne Lorenzo, Clare Murray-Lawson, Stephanie Shiers, Muhammad Saad Yousuf, Annemarie Dedek, Eve C. Tsai, Erin Vines, Peter Horton, Anna Cervantes, Tariq Khan, Geoffrey Funk, Gregory Dussor, Antoine G. Godin, Francesco Ferrini, Yves De Koninck, Michael E. Hildebrand, Theodore J. Price

## Abstract

Loss of GABAergic and glycinergic inhibitory efficacy in the spinal dorsal horn is associated with neuropathic pain and opioid-induced hyperalgesia in rodent models. Downregulation of the KCC2 chloride extrusion transporter is a key mechanism underlying this decreased inhibitory efficacy, but to-date there is no evidence supporting or opposing this hypothesis in humans. Here we demonstrate that KCC2 expression is decreased in superficial dorsal horn neurons of organ donors who died with a documented history of pain, or of long-term opioid use. We show profoundly decreased KCC2 dorsal horn membrane expression in a primary cohort associated with either chronic pain or opioid use, and in a replication cohort of mixed chronic pain and opioid use history. These results show that decreased dorsal horn inhibitory efficacy likely promotes chronic pain in humans and support the development of therapeutics augmenting KCC2 function as a treatment for chronic pain and opioid use disorders.

## Main text

Maintaining a balance of neuronal excitation and inhibition is fundamentally important for all neuronal circuits. Fast inhibition is mediated by GABA and glycine actions on ionotropic receptors that bind these neurotransmitters, GABA_A_ and glycine receptors. Opening of these ion channels allows for Cl^−^ flux through these channels, although the direction and magnitude of this anionic gradient is dependent on intracellular Cl^−^ concentration. In adult neurons, this gradient is largely dependent on the expression, localization, and activity of the K^+^, Cl^−^ cotransporter KCC2 ^1,2^. Efficient inhibition via GABA_A_ and glycine receptor ion channels requires low intracellular Cl^−^ concentrations which are maintained by plasma membrane-localized KCC2 cotransporters ^1,3,4^.

More than two decades of work in animal models demonstrate that tissue or nerve injury in the periphery causes a loss of inhibitory efficacy in the outer laminae of the dorsal horn of the spinal cord ^5^. This promotes pain and causes hypersensitivity to mechanical stimulation as inputs from dorsal root ganglion neurons to the spinal cord are not efficiently filtered, resulting in increased excitatory inputs to projection neurons that ultimately send nociceptive information to the brain ^1,5^. While several mechanisms to explain this loss of inhibitory efficacy have been proposed, decreased KCC2 expression, membrane localization, and function have been consistently reported in a variety of mouse and rat models of chronic pain ^3,6-12^. Decreased KCC2 function in the dorsal horn has also been reported in rodent models of opioid use disorder (OUD) suggesting that this mechanism may explain paradoxical pain caused by opioids, and some of the negative reinforcing effects of these drugs ^3,13,14^. Importantly, drugs that increase KCC2 function alleviate mechanical pain hypersensitivity in rodent chronic pain and OUD models ^3,13^.

A key gap in knowledge is whether KCC2 is downregulated in the spinal dorsal horn in humans with either chronic pain or OUD. Previous work on human spinal cord slices prepared from tissue recovered from organ donors demonstrates that brain derived neurotrophic factor (BDNF) can decrease membrane expression of KCC2, similar to findings in rodents^7^, but this effect was only seen in spinal cord slices prepared from male organ donors ^15^. Hence, while a key, sexual-dimorphic mechanism of KCC2 downregulation appears to be conserved in humans, the hypothesis that KCC2 is downregulated *in vivo* in humans with chronic pain or OUD remains untested. We sought to test this hypothesis directly using tissues recovered from organ donors with extensively documented medical histories.

From organ donor spinal cords recovered and preserved over the course of 4 years within North Texas in collaboration with the Southwest Transplant Alliance, we selected 18 lumbar spinal cords for the primary cohort for this study. All tissues were recovered within 1-3 hours of cross clamp and fresh frozen in the operating room using powdered dry ice ^16^. Tissues chosen for analysis showed no signs of damage to the dorsal horns from cause of death, or during dissection or the freezing process, which was confirmed using Hematoxylin and Eosin staining (**Figure 1**).

**Figure 1:**
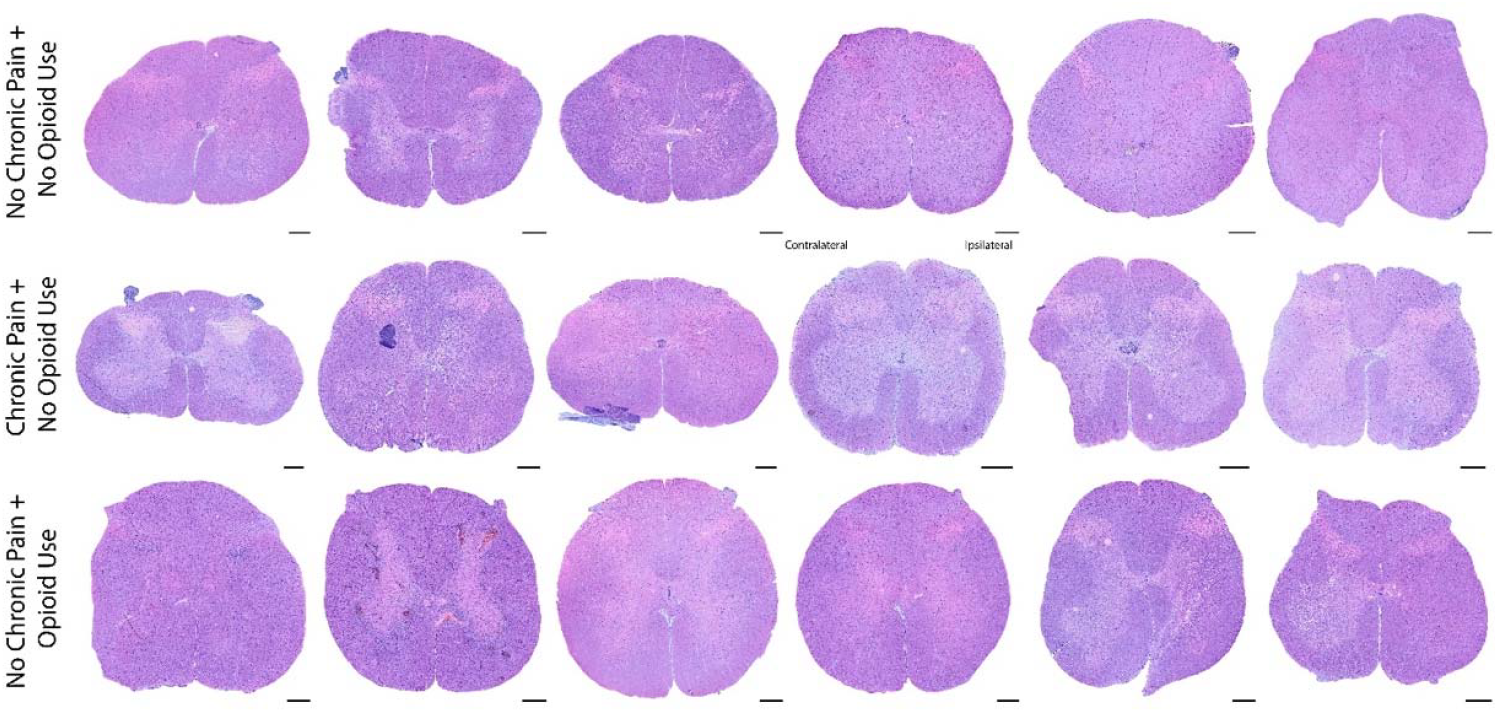
Hematoxylin & Eosin staining was conducted to confirm high quality tissue with no ice damage or trauma to the lumbar spinal dorsal horn in all organ donors prior to further analysis. Top row shows representative tissue from six donors with no chronic pain or opioid use, middle row is from six donors with medically defined chronic pain conditions and no opioid use, bottom row is from six organ donors with no history of chronic pain but a history of illicit opioid use (scale = 1mm; x20 magnification).

The primary cohort was comprised of 3 groups: 6 control donors with no known history of pain or drug use, 6 with a clear history of chronic pain affecting tissues with input to the lumbar spinal cord (only 1 having a history of prescribed opioid medication for pain treatment), and 6 donors with a history of illicit opioid use and a cause of death of drug overdose suggesting the presence of OUD (**Table 1**). The donors chosen were sex- and age-matched within all three groups and ranged in age from 30 to 60 years, with an ethnic diversity consistent with the population of the North Texas area (**Table 1**). We examined KCC2 immunoreactivity in at least 3 lumbar spinal cord sections from each organ donor using a previously described method ^17^ (see online methods) that employs NeuN immunolabelling to reveal cell bodies (**Figure 2**).

**Table 1:** Donor demographics of tissues used in the primary cohort processed at UTDallas.

| Donor ID | Age (yrs) | Sex | Ethnicity | Cause of Death | PMI (hrs) | Medical Conditions | Illicit Drug Use | Prescribed Medications | Classification | NDD vs DCD* |
| --- | --- | --- | --- | --- | --- | --- | --- | --- | --- | --- |
| UTD-DN0123 | 33 | Male | White | Head Trauma/ MVA | 2 | Nothing of note. | Nothing of note. | Nothing of note. | No Chronic Pain, No Opioid Use | NDD |
| UTD-DN0294 | 50 | Male | White/ Hispanic | CVA/ Stroke | 2 | Nothing of note. | Nothing of note. | Hypertension and diabetes medications. | No Chronic Pain, No Opioid Use | DCD |
| UTD-DN0043 | 63 | Male | White | Head Trauma/ Blunt Injury | 2 | Nothing of note. | Nothing of note. | Nothing of note. | No Chronic Pain, No Opioid Use | NDD |
| UTD-DN0158 | 36 | Female | Black | Anoxia/ Seizure | 1.5 | Nothing of note. | Nothing of note. | Hypertension and diabetes medications. | No Chronic Pain, No Opioid Use | NDD |
| UTD-DN0059 | 50 | Female | White | Head Trauma/ GSW | 2 | Nothing of note. | Nothing of note. | Nothing of note. | No Chronic Pain, No Opioid Use | NDD |
| UTD-DN0070 | 64 | Female | Black | CVA/ Stroke | 1.5 | Nothing of note. | Nothing of note. | Nothing of note. | No Chronic Pain, No Opioid Use | DCD |
| UTD-DN0095 | 34 | Male | Black | Anoxia/ Cardiac Arrest | 1.5 | Neuropathy in feet & back pain. | Marijuana smoked every few days for last 10 years. | Hypertension and diabetes medications, Gabapentin & Tylenol for pain. | Chronic Pain, No Opioid Use | NDD |
| UTD-DN0275 | 54 | Male | Black | Anoxia/ Natural Causes | 2.5 | Chronic abdomen pain / Crohn's disease, arthritis (knee and spine), migraine. | Nothing of note. | Antidepressants, Skyrizi infusions and Welchol for Crohn's, Oxycodone & Lyrica for pain. | Chronic Pain, No Opioid Use** | NDD |
| UTD-DN0129 | 60 | Male | White | Anoxia/ Cardiovascular | 3 | Peripheral neuropathy with frequent falls. | Nothing of note. | Gabapentin for pain. | Chronic Pain, No Opioid Use | DCD |
| UTD-DN0157 | 68 | Male | White | CVA/ Stroke | 1 | Left AKA due to painful diabetic neuropathy complications. | Historic cocaine and methamphetamine user > 10 years clean. | Diabetes medications. | Chronic Pain, No Opioid Use | NDD |
| UTD-DN0149 | 63 | Female | White | Anoxia/ Seizures | 1.5 | Fibromyalgia and osteoarthritis. | Nothing of note. | Antidepressants, anxiety, hypertension, seizure, anti-swelling, sleep and heartburn medications. Gabapentin & meloxicam for pain. | Chronic Pain, No Opioid Use | DCD |
| UTD-DN0225 | 63 | Female | White | Anoxia/ ICH Stroke | 2 | Osteoarthritis / knee pain. | Amphetamines used occasionally > 40 years ago. | Several medications including analgesics. | Chronic Pain, No Opioid Use | NDD |
| UTD-DN0143 | 31 | Male | Hispanic | Anoxia/ Drug Overdose | 2 | Nothing of note. | Hydrocodone, Marijuana, Xanax, Cocaine. | Diabetes medication. | No Chronic Pain, Opioid Use | NDD |
| UTD-DN0100 | 37 | Male | White | Anoxia/ Drug Overdose | 1 | Nothing of note. | Heroin, oxycodone laced with fentanyl, Cocaine, Benzodiazepines, Ambien | Nothing of note. | No Chronic Pain, Opioid Use | NDD |
| UTD-DN0183 | 39 | Male | White | Anoxia/ Drug Overdose | 2.5 | Nothing of note. | Cocaine, Methamphetamine, Toxicology report positive for opioids. | Asthma and cold sore medication. | No Chronic Pain, Opioid Use | NDD |
| UTD-DN0025 | 30 | Female | White | Anoxia/ Drug Overdose | 1 | Nothing of note. | Heroin, Methamphetamine, THC. | Nothing of note. | No Chronic Pain, Opioid Use | NDD |
| UTD-DN0108 | 36 | Female | White | Anoxia/ Drug Overdose | 2.5 | Nothing of note. | Toxicology report positive for THC, Opiates & Tylenol. | Nothing of note. | No Chronic Pain, Opioid Use | NDD |
| UTD-DN0119 | 50 | Female | Hispanic | Anoxia/ Drug Overdose | 2.5 | Nothing of note. | Heroin, Marijuana, Cocaine. | Nothing of note. | No Chronic Pain, Opioid Use | NDD |

\* Neurological Determination of Death vs Donation upon Circulatory Death.
\*\* Included in the Chronic Pain, No Opioid Use group as opioids were prescribed and not illicitly used, indicative of opioid-use disorder.

**Figure 2:**
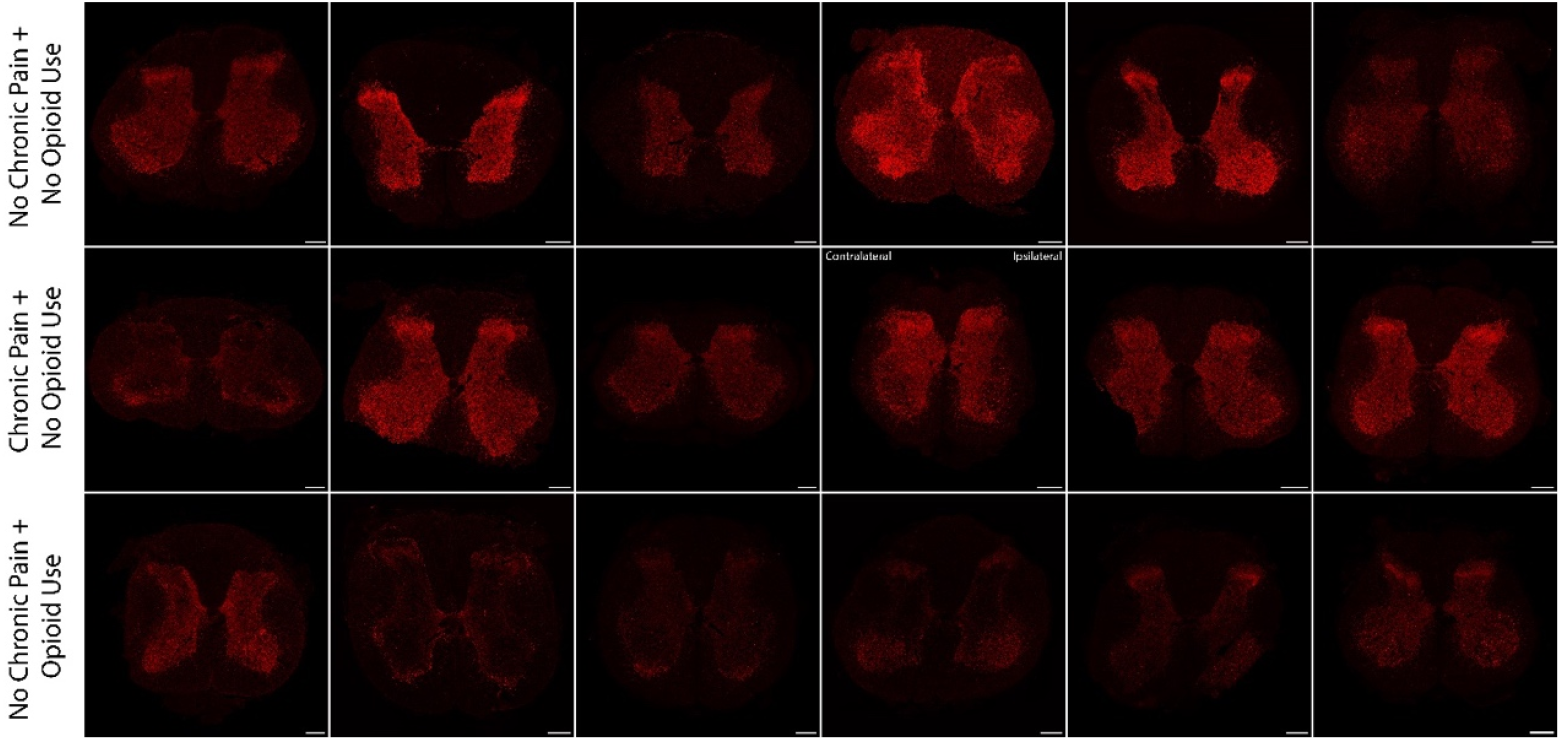
KCC2 immunolabelling in spinal cord sections from all organ donors in the primary cohort reveals staining within the grey matter with a dense plexus in lamina II. Top row shows representative tissue from six donors with no chronic pain or opioid use, middle row is from six donors with medically defined chronic pain conditions and no opioid use, bottom row is from six organ donors with no history of chronic pain but a history of illicit opioid use. All images were scanned with the same laser settings (scale bar = 1mm, x20 magnification).

We quantified membrane-associated KCC2 in laminae I and II dorsal horn neurons from at least 3 scans per lumbar spinal cord section from all organ donors, sampling at least 99 neurons per dorsal horn, with an average of 379 neurons per donor dorsal horn (representative images from each donor shown in **Figure 3** for lamina I and **Figure 4** for lamina II).

**Figure 3:**
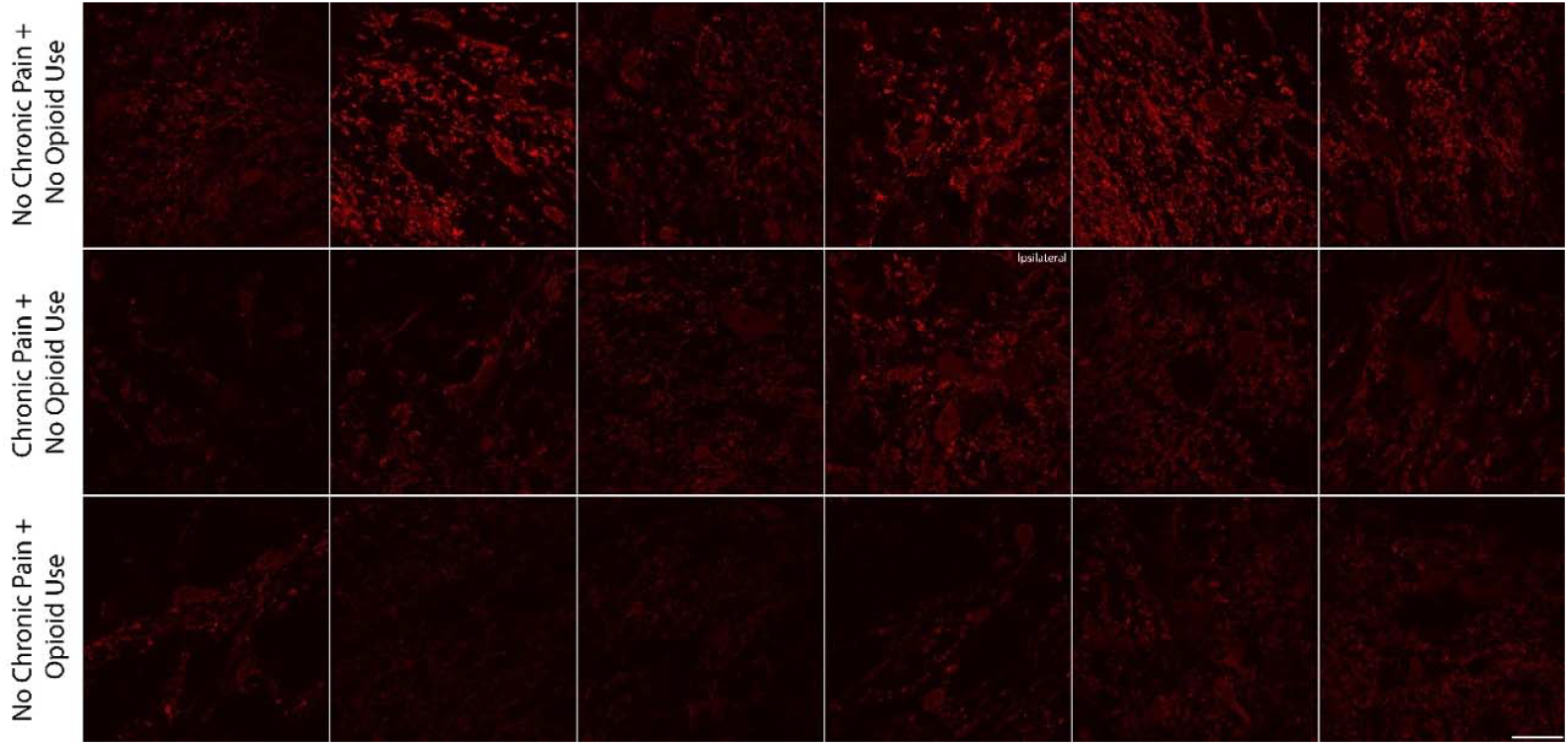
At a high magnification of lamina I, KCC2 immunoreactivity reveals dendritic and somatic membrane labelling, with some examples of internalization within the cell body. Representative images from all organ donors in the primary cohort used for analysis; top row shows representative tissue from six donors with no chronic pain or opioid use; middle row is from six donors with medically defined chronic pain conditions and no opioid use; bottom row is from six organ donors with no history of chronic pain but a history of illicit opioid use (all images to the same scale and confocal settings, scale bar = 50µm, x100 magnification).

**Figure 4:**
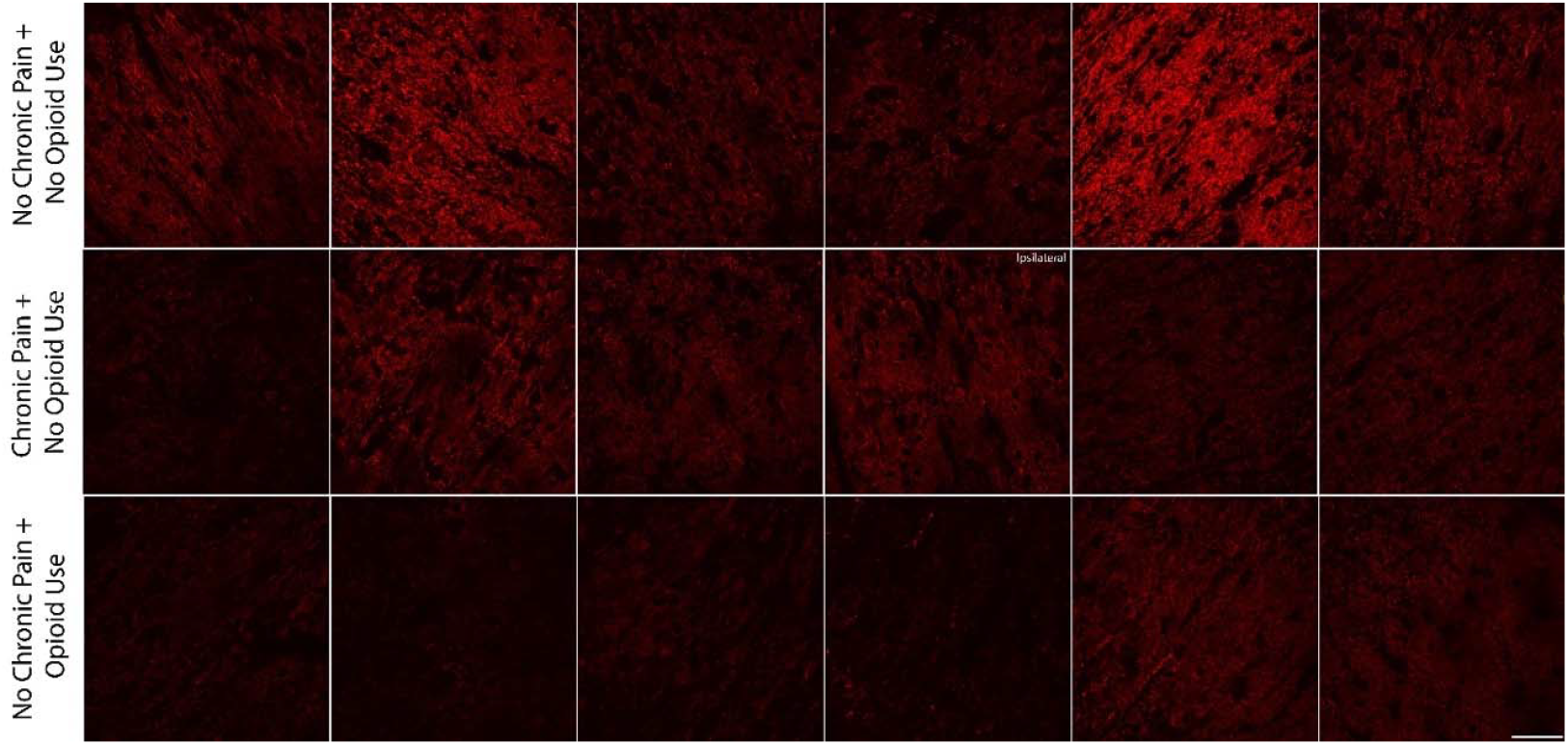
High magnification of KCC2 immunostaining in lamina II from all organ donors in the primary cohort reveals much denser dendritic and somatic membrane labelling than other areas of the spinal cord. Representative images from all organ donors in the primary cohort used for analysis: top row shows representative tissue from six donors with no chronic pain or opioid use; middle row is from six donors with medically defined chronic pain conditions and no opioid use; bottom row is from six organ donors with no history of chronic pain but a history of illicit opioid use. (all images to the same scale and using the same confocal settings, scale bar = 50µm, x100 magnification).

We found that membrane KCC2 was reduced in organ donors with a history of chronic pain (adjusted p value = 0.049) and in organ donors with a history of opioid use (adjusted p value = 0.0037) compared to control donors with no history of pain or opioid use (**Figure 5a-d**). There was no clear difference in membrane KCC2 immunoreactivity and any particular pain type (**Figure 5e**) consistent with animal model data where KCC2 downregulation has been reported in neuropathic pain ^6,8-11^, arthritis ^12^, visceral ^18^, and widespread pain models ^19^.

**Figure 5:**
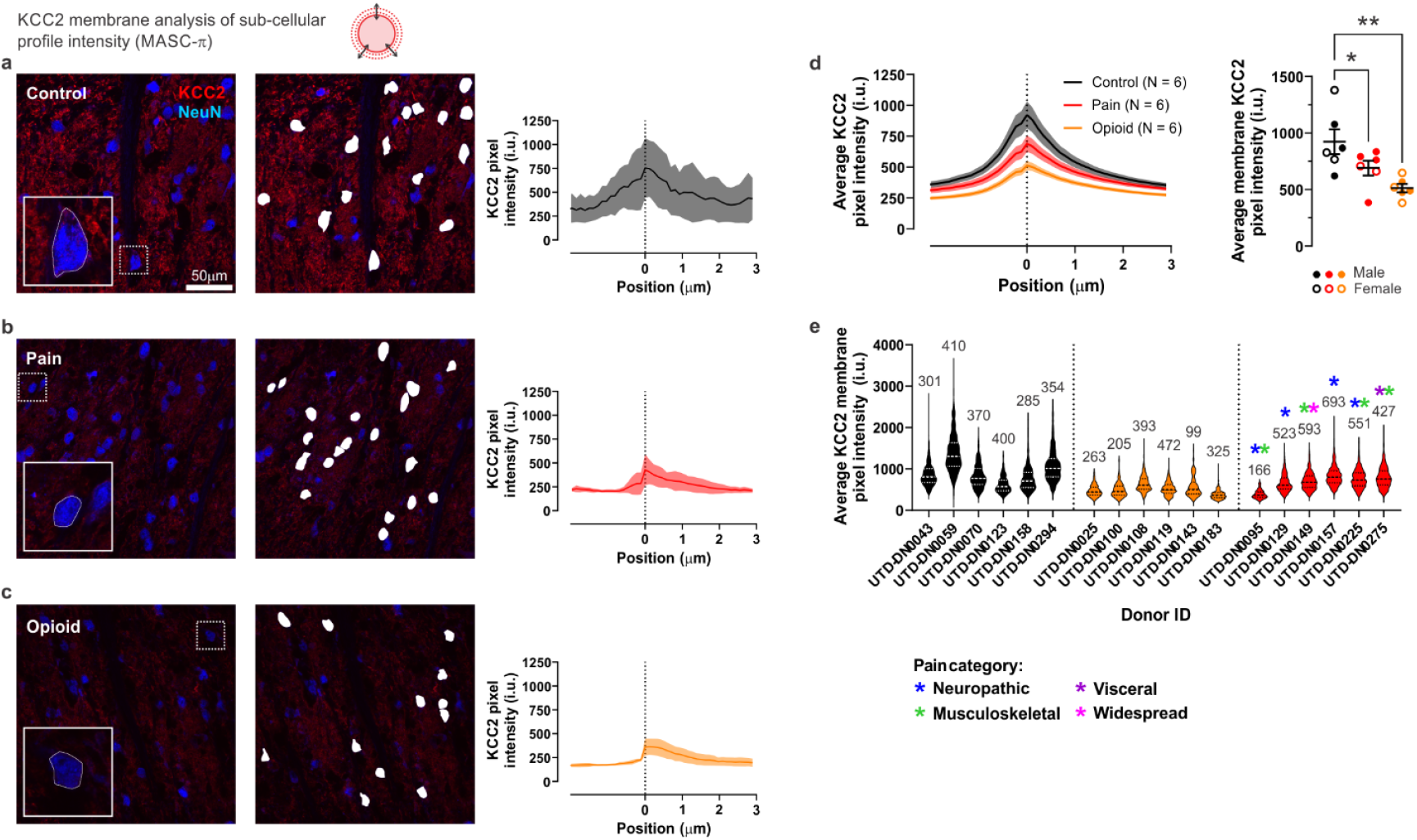
KCC2 membrane downregulation associated with pain or opioid use in primary cohort. Representative confocal images of NeuN and KCC2 from the superficial dorsal horn of (a) control, (b) chronic pain, and (c) opioid use donors with neuronal outlines (white). On the right, an example analysis of a single neuron for each group. (d) KCC2 intensity profiles at different positions along the membrane (left), with averaged membrane KCC2 for each group (right). (e) Violin plots of the KCC2 membrane intensity of all analyzed neurons separated by donor and group. Pain categories of chronic pain patients are represented with colored asterisks. Data are shown as mean ± S.E.M. \**P* < 0.05, \*\**P* < 0.01; one-way ANOVA with Holm-Sidak’s multiple comparison test.

Interestingly, in one organ donor sample with a medical history of a unilateral lower leg amputation there was a decrease in membrane KCC2 in the dorsal horn on the side ipsilateral to the amputation (**Figure 6**). While the sample size is too small for statistical analysis, there was also a clear trend for decreased membrane KCC2 in both male and female organ donors when separated by sex with either a history of chronic pain or opioid use (**Figure 7**), again consistent with animal model findings ^9^.

**Figure 6:**
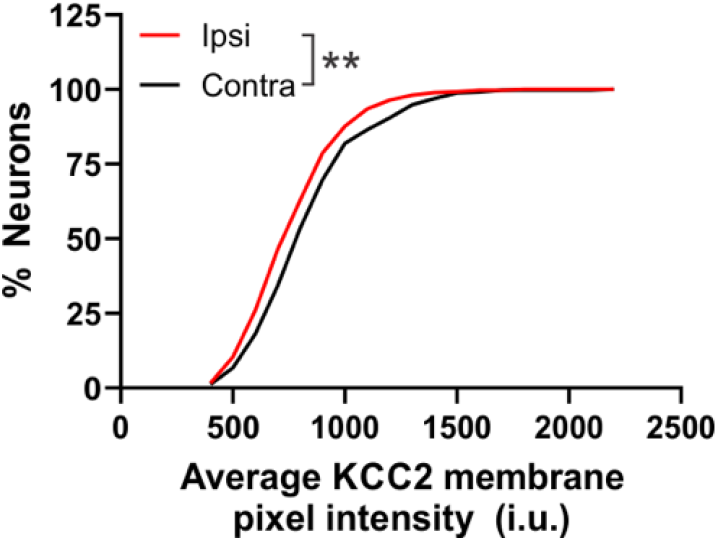
KCC2 membrane immunoreactivity downregulation in the dorsal horn ipsilateral to lower leg amputation. Cumulative frequency of average membrane KCC2 per neuron for donor UTD-DN-0157, separated by ipsilateral (ipsi) or contralateral (contra) side to injury.

**Figure 7:**
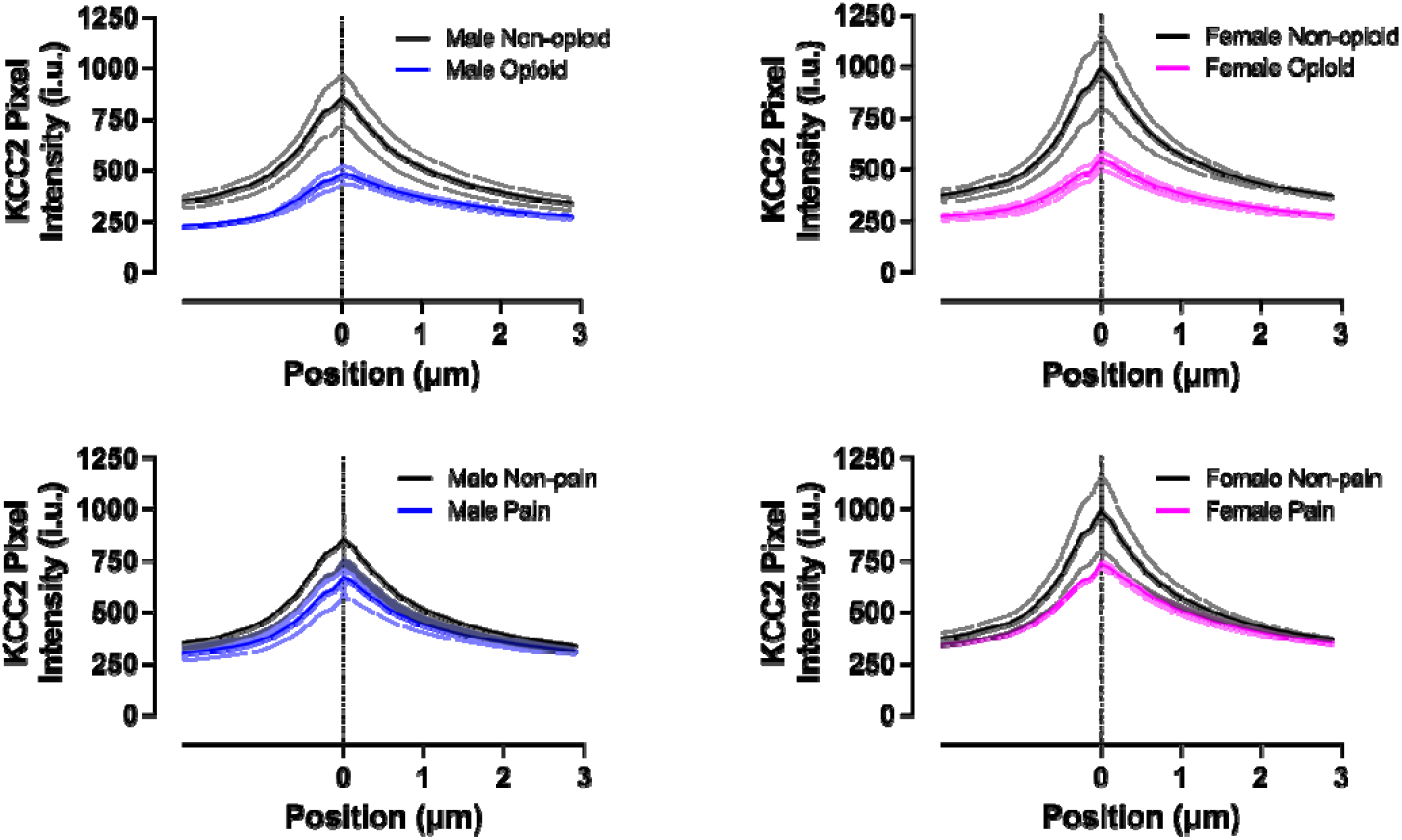
KCC2 membrane immunoreactivity per donor, separated by sex from the primary cohort. A limitation of the data generated from the primary cohort is the small sample size, however, we focused on donors with clear medical histories that allowed for classifying donors into distinct categories of no chronic pain or opioid use history, chronic pain, or opioid use. To increase the rigor of these findings we examined KCC2 membrane expression in the dorsal horn in a second cohort of organ donor spinal cords recovered at the Ottawa Hospital Research Institute and Carleton University in Ontario, Canada. These spinal cords were prepared for slice electrophysiology and/or biochemical assessment ^7,15^ and frozen prior to KCC2 immunostaining as described in online methods. These organ donor tissues could not be divided into pain-free, chronic pain, and opioid use history groups, but could be divided into two groups with no history of pain or opioid use and another with both a history of pain and opioid use (**Table 2**). Given the overall size of the donor pool in the replication cohort this overlap is expected given the widespread prescription for opioids within the chronic pain population in North America ^20^. In this replication cohort, we also found a significant decrease in membrane KCC2 in lamina I and II neurons in the pain and opioid use group (**Figure 8a-c**).

**Figure 8:**
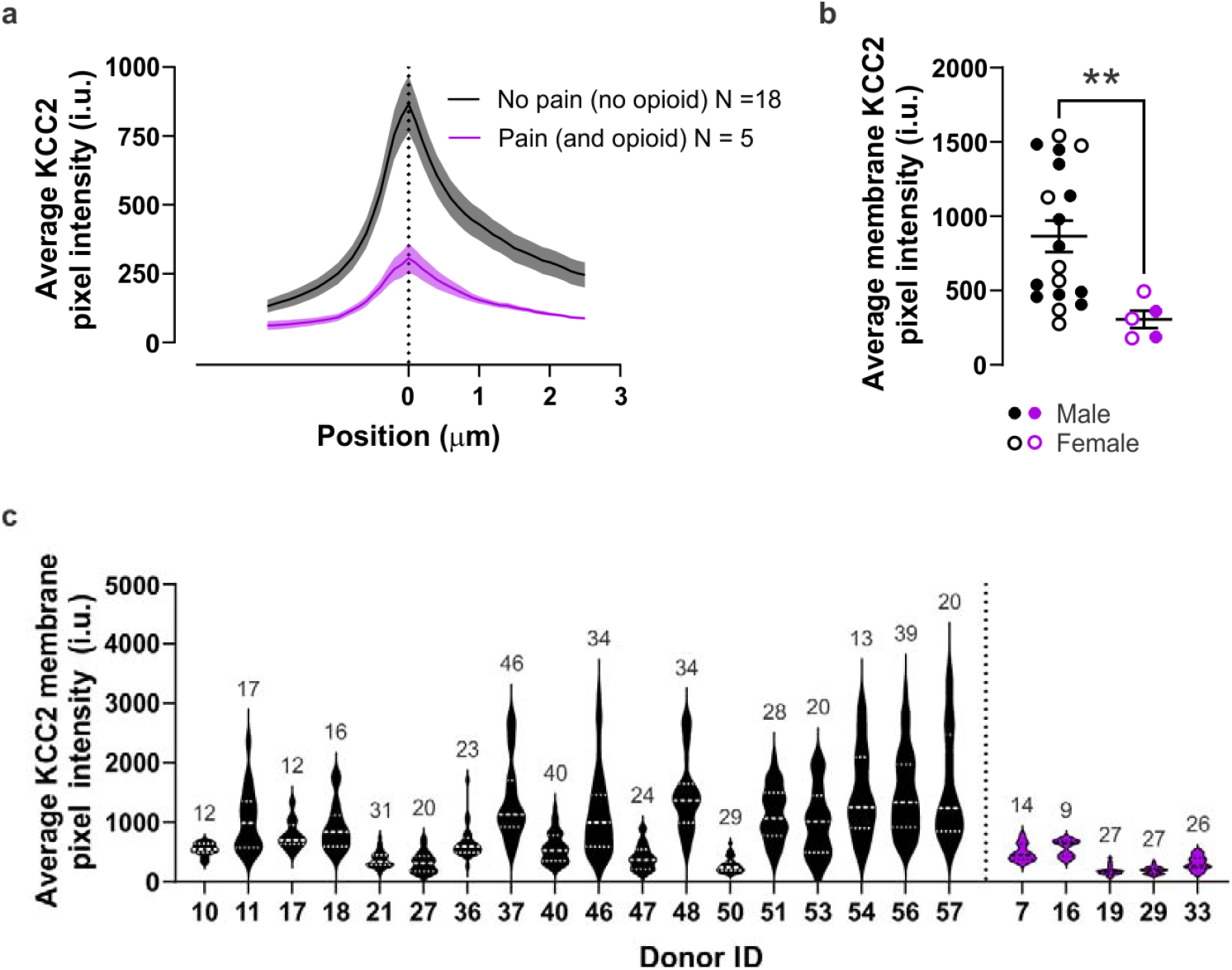
KCC2 membrane downregulation associated with pain and opioid use in replication cohort. (a) KCC2 intensity profiles at different positions along the membrane, with averaged membrane KCC2 for each group (b). (c) Violin plots of the KCC2 membrane intensity of all analyzed neurons separated by donor and group. Data are shown as mean ± S.E.M. \*\**P* < 0.01; Mann-Whitney test.

Kolmogorov-Smirnov test to compare cumulative distributions (p = 0.0034, n = 366 neurons for ipsi, n = 327 for contra).

**Table 2:** Donor demographics of tissues used in the replication cohort processed at Carleton University.

| Group | Age range (yrs) | Sexes | CODs | Medical Conditions | Illicit Drug Use | Prescribed Medications | NDD vs DCD |
| --- | --- | --- | --- | --- | --- | --- | --- |
| No Opioids and No Pain (n=18) | 18-70 | 11 males, 7 females | Hemorrhage (8), Anoxic brain injury (3), elevated intracranial pressure (3), TBI (2), stroke (1), MVA (1) | Nothing of note. | No (16), Yes (2) | Hypertension medications/ blood thinners (5), SSRIs (3) | NDD (15), DCD (3) |
| Opioids and Pain (n=5) | 20-70 | 2 males, 3 females | Hemorrhage (2), anoxic brain injury (1), cerebral thrombosis (1), meningitis (1) | Chronic back pain (3), chronic abdominal pain (1), Crohn's disease (1), migraines (1), fibromyalgia (1), osteoarthritis (1) | No (5) | Hydromorphone (3), gabapentin (1), oxycodone (1), Tylenol 3's (1), Celebrex (1), SSRIs (1) | NDD (5) |

Our work provides compelling evidence of KCC2 downregulation in the membrane of lamina I and II neurons of the spinal dorsal horn in humans with a history of chronic pain, a history of opioid use, or a combined history of both. Given the critical function of membrane KCC2 in maintaining low intracellular Cl^-^ concentration in neurons ^1,5,6^, this finding is consistent with decreased inhibitory efficacy in human lamina I and II neurons in people with chronic pain or OUD. A strength of our study is the replication of our key finding in two separate cohorts at geographically and demographically distinct sites. Whilst the second cohort provides only data on combined chronic pain and opioid use, the results offer mechanistic evidence in humans for the poor efficacy of opioids in relieving chronic pain symptoms given that opioids themselves trigger signaling cascades to downregulate KCC2. We acknowledge several limitations. First, the sample size in each individual cohort is small, but consistent with sample sizes in animal studies examining this target ^6,8,9,11,12,14,18,19^. Second, we have only examined downregulation of membrane localization of KCC2 and not total protein or RNA downregulation. Loss of KCC2 protein or RNA expression could have important implications for therapeutic targeting of KCC2 in pain or OUD ^3,13^. However, membrane localization is where KCC2 plays its most important role as a cotransporter in maintaining chloride gradient and thus neuronal inhibitory potential. Membrane-exclusive KCC2 cannot be evaluated by total protein or total RNA screening measurements. Finally, we have not identified the neuronal populations where KCC2 membrane localization is downregulated in human laminae I and II. Achieving this level of precision will require developing protein or mRNA markers for human dorsal horn neurons. In closing, our findings provide new insight into the underlying neurobiology of chronic pain and OUD in humans and support the continued development of positive modulators of KCC2 to treat human disease.

## Methods

### Tissue Procurement, Processing and Scanning

#### UTDallas

All human tissue procurement procedures were approved by the Institutional Review Board at the University of Texas at Dallas and collected in collaboration with the Southwest Transplant Alliance. Briefly, the caudal portion of the spinal cord was surgically dissected from human organ donors within 1-3 hours of cross clamp and immediately frozen using crushed dry ice, then stored at -80°C until use (see Supplementary Table 1 for donor information from UTDallas). Blocks from the lumbar region were embedded in OCT then cut into 20 µm sections using a cryostat and mounted onto charged slides. Sections from each donor were processed for Hematoxylin and Eosin staining. Briefly, slides were placed back into the -80°C freezer overnight then were immediately incubated in a 37°C oven for 1 minute to defrost. Slides were then placed in pre-chilled 100% methanol (Fisher Scientific, cat# 11367996) and left at -20°C for 30 minutes. Following this, excess methanol was removed by tapping the slide gently onto tissue and then sections were covered with 100% isopropanol (Fisher Scientific, cat# 1003269) and left for 1 minute. Once the excess was removed, the sections were allowed to air-dry for 7 minutes then were incubated with Mayer’s Hematoxylin solution (Agilent, cat# S330930-2) for a further 7 minutes. Slides were rinsed 3 times and then covered with Dako Bluing buffer (Agilent, cat# CS70230-2) for 2 minutes. After a further rinse, Eosin solution (Sigma Aldrich, Cat# HT110216) was added to each slide and left for 1 minute before a final 2 rinses. Slides were briefly air-dried for 2 minutes at room temperature then placed in the 37°C oven for 5 minutes to dry completely. Coverslips were applied using glycerol (Sigma Aldrich, cat# G9012) as the mounting medium and secured with nail polish. Sections were then scanned with an Olympus VS200 slide scanner and inspected to ensure a high tissue quality before proceeding.

Subsequently, three sections (>60 µm apart in the rostrocaudal axis) were processed for immunostaining and analysed per donor. The sections were fixed by submersion in 4% paraformaldehyde for 15 minutes, then dehydrated using sequential concentrations of ethanol (50, 70, 100, 100%) for 5 minutes each. After the final rinse, slides were allowed to air-dry and then a barrier was drawn using a hydrophobic marker. Sections were rehydrated in 0.1M phosphate buffered saline (PBS), then rinsed a further two times using fresh PBS for 10 minutes. The tissue was then incubated in a primary antibody cocktail of rabbit anti-KCC2 (1:250; Sigma, Cat# 07-432), mouse anti-NeuN (1:500, Millipore, Cat#MAB377) and rat anti-substance P (1:100; Biorad, Cat# 8450-0505) in PBS with 0.3% triton and 5% normal goat serum, and left overnight at 4°C. Following 3 further rinses of PBS, the sections were then incubated in species-specific secondary antibodies (1:500; Invitrogen; cat# A21428, A21240, PA1-84761) for another night at 4°C. Slides were rinsed and coverslipped using Vectashield mounting medium (Vector Laboratories, cat # H-1000-10). A secondary antibody control reaction was run in parallel for each donor by reacting in the same way as above but with no primary antibody during the initial incubation (not shown).

Tissue was scanned using an Olympus FV3000RS microscope with a x40 oil-immersion lens and a 1.34x zoom (optimal for the aperture and lens used) to reach a resolution of 0.12µm per pixel. The laser settings and scanning conditions were kept the same between all donors. At least 3 scans per spinal section were taken from laminae I and II of both dorsal horns. The boundaries of laminae I and II were defined using a combination of brightfield illumination, the density of NeuN immunolabelling and the plexus of substance P immunolabelling (although this was not scanned or analysed). Secondary controls were scanned but not analysed.

#### UCarleton & ULaval

All human spinal tissue from adults was collected from organ donors identified through the Trillium Gift of Life Network. Ethics approval was obtained from the Ottawa Health Science Network Research Ethics Board (Protocol ID # 20150544-01H) and the Carleton University Research Ethics B Committee (Ethics Protocol #104836) to collect and conduct experiments with human tissue. The ethics protocol of this secondary cohort detail that multiple identifiers cannot be disclosed for any *individual* donor. For this reason, Supplementary Table 2 details the demographic information within each group as opposed to detailed summaries of each donor. All human tissue donors were screened for any potential blood-borne illnesses such as HIV, hepatitis, and syphilis. Additionally, information regarding significant illnesses and medications were collected if relevant to the present study, such as a previous diagnosis with a chronic pain condition, persistent use of pain-modulating drugs, or any other condition that might have contributed to spinal cord damage. Any donor with an incomplete medical history was excluded from analyses. Human donors were eligible if death determinations were made through neurological criteria (donation after neurological determination of death, NDD), or circulatory criteria (donation after circulatory death, DCD).

A sledge freezing Vibratome Leica VT1200S (Leica Microsystems) was used to cut 25 µm transverse sections of paraformaldehyde-fixed human spinal tissue. Sections were permeabilized in PBS (pH 7.4) with 0.2% Triton (PBS+T) for 10 minutes, washed twice in PBS, then incubated for 12 hours at 4°C in primary rabbit anti-KCC2 antibody (1:1000, Millipore/Upstate, Cat. #07–432), mouse anti-CGRP antibody (1:5000; Sigma #C7113) and chicken anti-NeuN antibody (1:1000, MilliporeSigma Cat. #6B9155) diluted in PBS+T containing 10% normal goat serum. Tissue was washed in PBS, and subsequently incubated for 2 hours at room temperature in goat-Cy3 anti-rabbit purified secondary antibody (1:500, Jackson ImmunoResearch Laboratories, Cat. #111–165–144) goat anti-chicken Alexa Fluor® 647 secondary antibody (1:500, Invitrogen Cat. #AB2535866) diluted in PBS+T (pH 7.4) containing 10% normal goat serum. Spinal sections were mounted on SuperFrost™ gelatin-subbed slides (Fisherbrand) and were cover-slipped using fluorescence mounting medium (Dako, Cat. #S3023).

A Zeiss LSM 880 Confocal Laser Scanning Microscope was used to acquire all confocal images. Acquisitions were 12-bit images, 2048 × 2048 pixels with a pixel dwell time of 10 µs. An oil-immersion ×63 plan-apochromatic objective was used for high magnification confocal laser scanning microscopy images, which were processed for quantification. Laser power was adequately chosen to avoid saturation and limit photobleaching. All acquisitions were performed using the same laser settings (laser, power, photomultiplier tube (PMT) settings, image size, pixel size and scanning time).

The superficial dorsal horn was defined as the area containing CGRP-immunoreactive primary afferent terminals. Only neurons present in this were scanned and considered for analysis. For both imaging and analysis, the experimenter was blinded to the experimental conditions.

#### Image Analysis & Statistics

The membrane analysis of sub-cellular profile intensity (MASC-_π_) was used as previously to quantify KCC2 levels at the membrane of neurons ^17^. The membrane KCC2 staining of lamina I and II NeuN+ cells was manually delineated on each image. The intracellular NeuN staining was used instead to delineate the cell border when KCC2 could not be discriminated from the background to avoid bias against cells with low KCC2. Cells that didn’t have a clear membrane KCC2 or a uniform NeuN staining were not considered for analysis. The analyser was blinded to each subject’s sex and group, and the brightness of each image was adjusted automatically during analysis to eliminate bias. The drawn region of interest was enlarged and reduced by ± 3 μm, and the mean KCC2 pixel intensity and standard deviation were calculated as a function of the distance to the drawn region of interest for each cell. Positive values correspond to the intracellular space whilst a negative position value represents the region outside of the labelled neuron. All cells of a given subject were averaged together, and each subject’s mean was used for statistical analysis.

## Acknowledgements

The authors thank the organ donors and their families for their gift of life.

